# Pydigree: a python library for manipulation and forward-time simulation and of genetic datasets

**DOI:** 10.1101/213413

**Authors:** James E. Hicks

## Abstract

The development of software for working with data from population genetics or genetic epidemiology often requires substantial time spent implementing common procedures. Pydigree is a cross-platform Python 3 library that contains efficient, user friendly implementations for many of these common functions, and support for input from common file formats. Developers can combine the functions and data structures to rapidly implement programs handling genetic data. Pydigree presents a useful environment for development of applications for genetic data or rapid prototyping before reimplementation in a higher-performance language.

Pydigree is freely available under an open source license. Stable sources can be found in the Python Package Index at https://pypi.python.org/pypi/pydigree/, and development sources can be downloaded at https://github.com/jameshicks/pydigree/

## 1 Introduction

Development of applications and algorithms for genetic datasets requires reimplementing many basic functions and parsing common file formats. Identifying bugs and accounting for edge-cases represents a substantial use of developer time. Genetic datasets are often very large, and inefficient processing can result in lost productivity. A common implementation of these functions would allow for faster development time, fewer bugs,. Pydigree is a python framework for genetic datasets that attempts to solve this problem.

The Python programming language is commonly used in bioinformatics, and users have access to a wide range of libraries for numerical computing, statistics, and machine learning. While there are python packages that work with genetic data, such as simuPOP [8], they tend to focus on simulation or specialized functions. Pydigree provides a general purpose python module containing implementations of common procedures on datasets of diploid subjects. This allows developers to integrate their genetics algorithms and simulations with the other python libraries.

## 2 Features

Pydigree provides a ‘pythonic’ environment for creation and manipulation of genetic data. Entities and common procedures are implemented in an object-oriented fashion, with a focus on organized selection and manipulation of data. Where possible, functions are implemented lazily (deferring computation until the result is required). For common computationally intensive tasks, functions have been implemented in the C++ or Cython [1] programming languages. The module provides these data structures and functions independently, allowing users to combine them as needed in their own programs.

Pydigree is designed to be extensible and easy to integrate with other programs. The python ecosystem contains many useful packages for scientific computation, and pydigree can translate its objects to data structures usable by them. The provided data structures can be used in python’s language interoperability features, allowing access to functions written in C/C++ or R.

Users will most commonly interact with three kinds of objects: the individuals themselves, collections of individuals, and genotypes. Individual represents the set of genotypes and phenotypes for an individual in the dataset. Population, Pedigree, and PedigreeCollection are collections of individuals. They provide convenient functions for selecting certain individuals and batch operations for each individual in the collection. Additionally, collections can calculate allele frequencies. Population represents a collection of unrelated individuals. Pedigree is a subclass of Population and represents a group of individuals connected by a known family structure. It provides common functions for pedigrees, including kinship and inbreeding coefficients. PedigreeCollection represents a collection of pedigree objects and can perform batch operations on them. When genealogies are present, they can be navigated through using the paths submodule.

The third kind of major object organizes genetic information. The main object here is Alleles, which carries a haploid chromosome’s set of alleles. The alleles are stored efficiently and can be queried and sliced like other python objects. For each chromosome in a population, there is a single ChromosomeTemplate object. This class organizes the information on each variant site in the chromosome, including allele frequencies and both physical and genetic positions. When handling data generated from sequencing experiments problems with memory usage can often arise. Pydigree provides a data structure, SparseAlleles, which stores only minor alleles. Since the bulk of variation in the human genome is rare, the sparse container can result in a substantial reduction in memory use for large datasets.

Pydigree supports file IO in a variety of formats, including VCF [2], PLINK, and BEAGLE files. GZIP, BZIP2 and LZMA/XZ compressed files are handled seamlessly. Pedigrees can be input in the popular LINKAGE format used by MERLIN and PLINK, and genotypes from a separate file may be merged in.

## 3 Simulation

Pydigree contains functions for the simulation of both pedigrees and population-based datasets by a forward-time strategy. An initial population or pool of chromosomes can be iterated forward in time by mating randomly chosen members and using their offspring for the next generation. Genealogical history for each individual in a generation can be retained. Existing datasets can be used as initial chromosomes for simulation, randomly generated.

When intermediate genotypes do not need to be realized (e.g. no effects of selection or assortative mating), datasets can be generated rapidly. Instead of genotypes, individuals carry references to segments on a chromosome in the initial generation. This has two main advantages. First, it reduces the amount of time is spent allocating and copying haplotypes between individuals. Second, true haplotype phase and identity-by-descent states are known through the simulation. This has a variety of possible uses, including simulating pedigrees constrained by inheritance patterns or testing methods for detecting IBD segments from genotypes. Once generation advancement is complete, genotypes are assigned to the founder generation and all other genotypes are resolved by their references to founder individuals.

### 3.1 Simulation of quantitative genetic models

Pydigree allows for simulated phenotypes to be generated from genotypes. Each allele can be given additive and dominance effects, collected in the class QuantitativeTraitArchitecture. At the end of simulation, this class combines the effects of all specified alleles to give the overall genetic effect. If a dichotomous trait is required, a threshold can be specified such that phenotypes above it are marked affected. The sum of the effects can be rescaled and combined with noise to give the trait a specified heritability. QuantitativeTraitArchitecture can optionally randomly select a set of markers or add extra dummy chromosomes to simulate the effect of polygenes on the trait. Additionally, this model can incorporate the effects of specified environmental exposures.

## 4 Included statistics and algorithms

Estimation of variance components is common in both genetic epidemiology and animal breeding. This is generally done via the linear mixed effects model (LMM). Pydigree implements an LMM that seamlessly integrates with the other objects. The MixedModel class can utilize outcome variables fixed predictors from collections of Individual objects. Random effects are specified by RandomEffect objects and allow for arbitrary incidence and covariance matrices. The model can be fit for maximum likelihood or restricted maximum likelihood using expectation-maximization [3] or Newton-like optimization methods. When using Newton-like methods, Newton-Raphson, Fisher Scoring [7], and Average-Information [4] methods are available. Math is done internally by the optimized linear algebra routines in the numpy [9] and scipy [5] software packages. To compare models a likelihood-ratio test is provided.

Identifying genome regions shared identical-by-descent (IBD) using genotype data is an increasingly common operation on populations. Pydigree provides a submodule, sgs, which uses the insight of Kong *et al.* [6], that IBD regions between a pair of individuals can not contain opposite homozygote genotype pairs. SGSAnalysis attempts to find these shared genome segments by calculating identity-by-state (IBS) across each chromosome and finds regions that do not have any IBS=0 genotype pairs. The algorithm used allows for user-defined acceptable rates of genotyping errors and genotype missingness.

Hidden Markov models (HMMs) are used in many algorithms for computational biology. Pydigree provides two implementations: a standard HMM, and a ‘genotype HMM’ which can vary its emission probabilities across observations (e.g. for alleles with different frequencies). Hidden states can be resolved by maximum likelihood (forwards-backwards algorithm) or maximum a posteriori (Viterbi decoding) methods.

## 5 Conclusion

Pydigree provides a useful and user-friendly development environment for genetic software. Pydigree also provides several programs as part of its installation. These scripts perform common tasks like kinship and inbreeding coefficient calculation (kinship.py), variance component estimation (polygenic.py), and forward time simulation of pedigree datasets (simulate pedigree data.py). While useful for their own purposes they also provide an example of how to use the pydigree library for common tasks.

Pydigree is available for download under an the open source Apache 2.0 license. Stable versions can be found at in the Python Package Index at https://pypi.python.org/pypi/pydigree/. Source code and developmental versions can be found at https://github.com/jameshicks/pydigree/.

## Acknowledgements

I would like to thank William Scott and Michael Province for their support during the development of this software.

## References

1. S. Behnel, R. Bradshaw, C. Citro, L. Dalcin, D. Seljebotn, and K. Smith. Cython: The best of both worlds. Computing in Science Engineering, 13(2):31–39, 2011.

2. P. Danecek, A. Auton, G. Abecasis, C. A. Albers, E. Banks, M. A. DePristo, R. E. Handsaker, G. Lunter, G. T. Marth, S. T. Sherry, G. McVean, R. Durbin, and. The variant call format and vcftools. Bioinformatics, 27(15):2156, 2011.

3. A. P. Dempster, N. M. Laird, and D. B. Rubin. Maximum likelihood from incomplete data via the em algorithm. JOURNAL OF THE ROYAL STATISTICAL SOCIETY, SERIES B, 39(1):1–38, 1977.

4. A. R. Gilmour, R. Thompson, and B. R. Cullis. Average information reml: An efficient algorithm for variance parameter estimation in linear mixed models. Biometrics, 51(4):1440–1450, 1995.

5. E. Jones, T. Oliphant, P. Peterson, et al. SciPy: Open source scientific tools for Python, 2001. [Online; accessed 2017-01-18].

6. A. Kong, G. Masson, M. L. Frigge, A. Gylfason, P. Zusmanovich, G. Thorleifsson, P. I. Olason, A. Ingason, S. Steinberg, T. Rafnar, P. Sulem, M. Mouy, F. Jonsson, U. Thorsteinsdottir, D. F. Gudbjartsson, H. Stefansson, and K. Stefansson. Detection of sharing by descent, long-range phasing and haplotype imputation. Nat. Genet., 40(9):1068–1075, Sep 2008.

7. K. Lange, J. Westlake, and M. A. Spence. Extensions to pedigree analysis. III. Variance components by the scoring method. Ann. Hum. Genet., 39(4):485–491, May 1976.

8. B. Peng and M. Kimmel. simuPOP: a forward-time population genetics simulation environment. Bioinformatics, 21(18):3686–3687, Sep 2005.

9. S. van der Walt, S. C. Colbert, and G. Varoquaux. The numpy array: A structure for efficient numerical computation. Computing in Science & Engineering, 13(2):22–30, 2011.

